# Deficient autophagy drives aging in *Hydra*

**DOI:** 10.1101/236638

**Authors:** Szymon Tomczyk, Quentin Schenkelaars, Nenad Suknovic, Yvan Wenger, Kazadi Ekundayo, Wanda Buzgariu, Christoph Bauer, Kathleen Fischer, Steven Austad, Brigitte Galliot

## Abstract

*Hydra* exhibits a negligible senescence as its epithelial and interstitial stem cell populations continuously divide. Here we identified two *H. oligactis* strains that respond differently to interstitial stem cell loss. Cold-resistant *(Ho_CR)* animals adapt and remain healthy while cold-sensitive *(Ho_CS)* ones die within three months, after their epithelial stem cells lose their selfrenewal potential. In *Ho_CS* but not in *Ho_CR* animals, the autophagy flux is deficient, characterized by a low induction upon starvation, proteasome inhibition or Rapamycin treatment, and a constitutively repressed Ulk activity. In the non-aging *Hydra vulgaris*, WIPI2 silencing suffices to induce aging. Rapamycin can delay aging by sustaining epithelial self-renewal and regeneration, although without enhancing the autophagy flux. Instead Rapamycin promotes engulfment in epithelial cells where p62/SQSTM1-positive phagocytic vacuoles accumulate. This study uncovers the importance of autophagy in the longevity of early-branched eumetazoans by maintaining stem cell renewal, and a novel anti-aging effect of Rapamycin via phagocytosis.

---

Studies using short-lived invertebrate organisms as the fruit fly or the nematode dramatically improved our understanding of aging (Longo and Finch 2003). However these model systems have some drawbacks such as developmental pausing, which might be activated when lifespan is experimentally prolonged, thus implying a mechanism different from mammalian aging (Austad 2009). Also except for the *Drosophila* gut, somatic tissues in adult flies and nematodes do not self-renew, while self-renewal is an essential component of homeostasis in humans. Finally, a significant proportion of human orthologous genes were lost in fly and nematodes as evidenced by their presence in cnidarians, a bilaterian sister group (Figure 1A) (Wenger and Galliot 2013; Schenkelaars et al. 2017). Therefore, additional invertebrate models could be profitably developed to help discover novel genes, pathways and mechanisms relevant for human aging.

**Figure 1:**
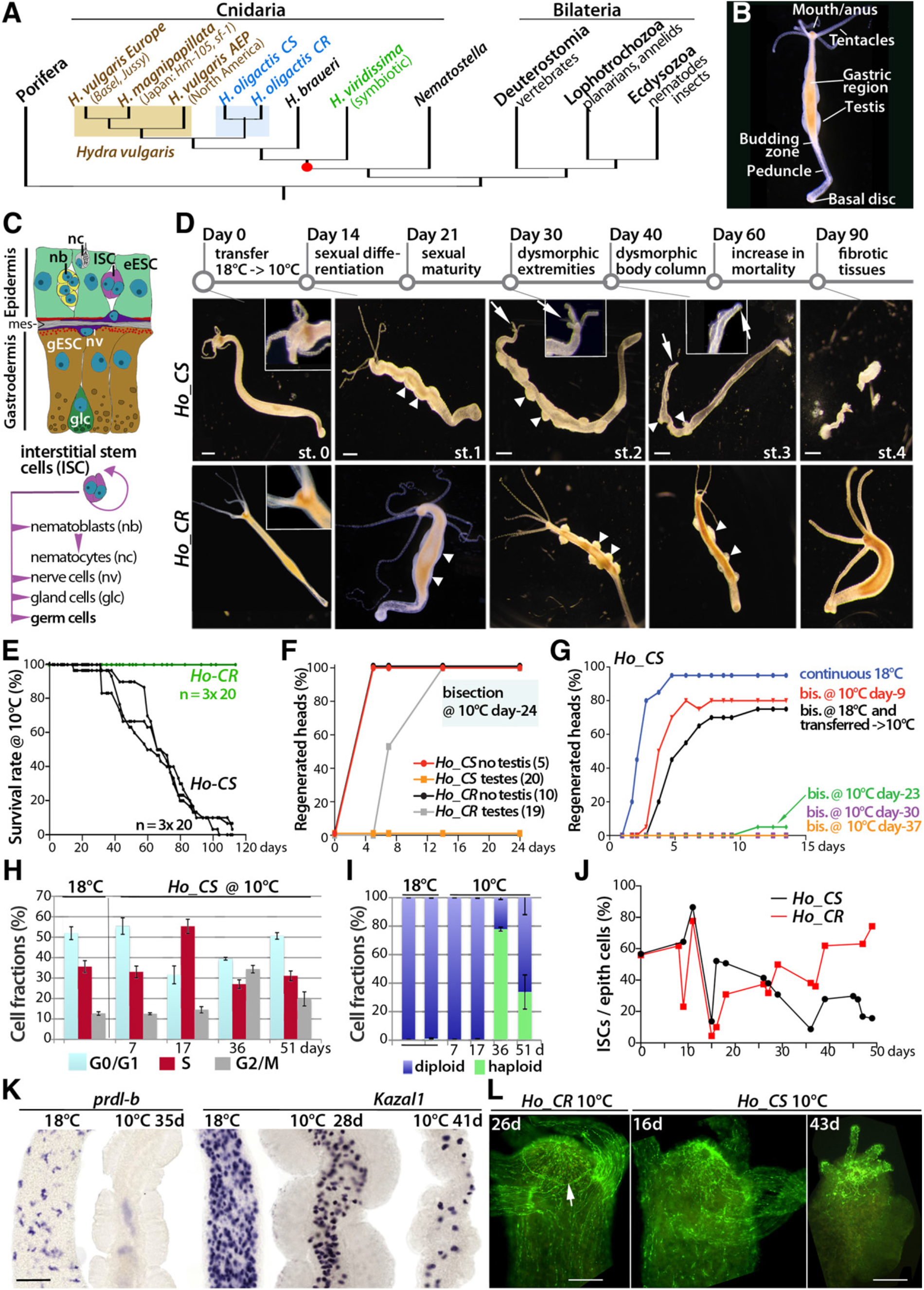
Inducible aging phenotype in cold sensitive *Hydra oligactis (Ho_CS)*. **(A)** Phylogenetic position of *Hydra* among metazoans. **(B)** Anatomy of a male *H. oligactis* animal. **(C)** Schematic view of *Hydra* gastric tissue; epidermal Epithelial Stem Cell (eESC), gastrodermal Epithelial Stem Cell (gESC), mesoglea (mes). (D) Morphological changes observed in ***Ho_CS*** (upper) and ***Ho_CR*** (lower) animals after transfer to 10°C (day 0); arrowheads: testes, arrows: degenerating head regions. Scale bar: 500 *μ*m. **(E)** Survival rates among ***Ho_CR*** and ***Ho_CS*** cohorts. **(F)** Head regeneration measured in ***Ho_CR*** or ***Ho_CS*** animals selected for the presence or absence of testes and bisected at mid-gastric level after 24 days at 10°C. **(G)** Head regeneration measured in ***Ho_CS*** animals bisected (bis.) either immediately before transfer to 10°C or 9, 23, 30 or 37 days after transfer. **(H, I)** Modulations in DNA content profiles detected by flow cytometry in ***Ho_CS*** animals maintained at 18°C or 10°C. **(J)** Ratio of ISCs (single and pairs) over epithelial cells counted in macerated tissues (triplicates of ~300 cells). **(K)** Progressive elimination in cold-exposed ***Ho_CS*** of the *prdl-b* and *Kazal1* expressing-cells that correspond to nematoblasts and gland cells respectively. Scale bar: 200 μm. **(L)** RFamide-positive neurons in apical regions of ***Ho_CR*** and ***Ho_CS*** animals; arrow: nerve ring. Scale bar: 100 μm.

*Hydra* is a small carnivorous freshwater cnidarian polyp (Figure 1B), characterized by a dynamic homeostasis and the ability to regenerate any missing part after amputation (Galliot 2012). It expresses a large number of vertebrate orthologs along its radially organized bilayered body plan. In culture conditions mortality of *Hydra vulgaris (Hv)* remains negligible over the years (Brien 1953; Martinez 1998) and well-fed animals remain asexual, reproducing by budding without showing replicative senescence (Schaible et al. 2015). Hence, *Hydra* seems to escape aging, an unusual property likely attributed to its three distinct stem cell populations that constantly self-renew in the central body column. The epidermal and gastrodermal epithelial stem cells (eESC, gESC) are multifunctional unipotent stem cells, while the interstitial stem cells (ISC) are multipotent, providing both somatic and germ cells (Figure 1C). ISCs and interstitial progenitors (collectively named i-cells) cycle much faster than ESCs; as a result animals transiently exposed to anti-proliferative drugs rapidly lose their i-cells and progressively become epithelial (Marcum and Campbell 1978; Sugiyama and Fujisawa 1978; Buzgariu et al. 2014). If force-fed, such epithelial *Hydra* remain viable, able to bud and regenerate (Marcum and Campbell 1978; Sugiyama and Fujisawa 1978), likely due to the rapid adaptation of their ESCs that modify their genetic program (Wenger et al. 2016).

However aging was reported in a species named *H. oligactis (Ho)* (Figure 1A) (Brien 1953; Littlefield et al. 1991). A rapid temperature drop to 10°C suffices to induce gametogenesis in *Ho* animals that rapidly stop budding, produce gametes and degenerate. This process, characterized by somatic i-cell loss, cytoskeleton disorganization, decline in body movements and feeding behavior, was identified as aging (Yoshida et al. 2006). In male and female *Ho* strains, sexual animals die within four months, showing Gompertzian mortality dynamics (Finch 1990). As *Ho* animals maintained at 18°C exhibit no signs of aging, we used cold transfer to induce aging experimentally in two *Ho* strains, one cold-sensitive *(Ho_CS)* that undergoes aging, and another, cold-resistant *(Ho_CR)* that survives gametogenesis. We show that these two strains exhibit striking differences in autophagy efficiency and in stem cell self-renewal. An inducible autophagy flux appears essential to prevent aging in *Hydra*, as *WIPI2* silencing suffices to shorten *Hv* survival. In *Ho_CS* a chronic Rapamycin treatment delays aging by stimulating epithelial phagocytosis and lipid droplet accumulation without rescuing autophagy.

## RESULTS

### *Ho_CR* and *Ho_CS* respond differently to cold exposure

To investigate aging in *Ho*, we used two closely related male strains that exhibit similar budding regulation at 18°C (_Figure_S1A_) but respond differently to cold exposure. After cold transfer, ~70% *Ho_CR* animals remain asexual and healthy while ~30% differentiate testes that reach maturity within 25 days, then lose sexual traits and return to physiological fitness without stopping budding or exhibiting aging signs (Figure 1D,1E, **Figure_S1**). After 300 days at 10°C, all *Ho_CR* animals are healthy, some showing mild dysmorphic signs (e.g. duplicated basal region, non-detached buds, not shown). By contrast, after transfer to 10°C, *Ho_CS* animals stop budding (**Figure_S1B**), 10% remain asexual, while 90% differentiate testes and show irreversible dysmorphic signs such as tentacle shrinking, head loss, stenosis of the body column as reported previously (Yoshida et al. 2006). Their survival time negatively correlates to the testis number (**Figure_S1D,E**). Within one month the animals lose the ability to regenerate (Figure 1F, 1G), show behavioral defects (**Movie-S1A, S1B**), while aging becomes irreversible (**Figure_S1H-O**). Hence *Ho_CS* but not *Ho_CR* animals undergo aging in response to cold-induced gametogenesis.

### Different cellular impact of gametogenesis in aging and non-aging *Hydra*

To monitor the impact of gametogenesis on somatic interstitial derivatives, we first analyzed DNA profiles by flow cytometry (Buzgariu et al. 2014) and found an increase in S-phase cells at day-17, when testes produce spermatogonia, and a large proportion of haploid cells at day-36 (Figure 1H,1I). Next we found the ratio of somatic i-cells over ESCs steadily decreasing in *Ho_CS* but only transiently altered in *Ho_CR* (Figure 1J), indicating that germ cell production leads to depletion of somatic i-cells in *Ho_CS*. Indeed, nematoblasts, gland cells, neurons disappear within few weeks after transfer to 10°C in *Ho_CS* (Figure 1K,1L) while persisting in *Ho_CR* even after 300 days at 10°C (not shown). Next we analyzed by RNA-seq the regulation of 20 i-cell markers in aging and non-aging *Ho* animals (**Figure_S2A**) and found several i-cell genes *(cnnos2, PaxA, vasa1, vasa2, ZNF845, cnox-2)* already up-regulated at day-25/day-32 in *Ho_CR* but not in *Ho_CS* (**Figure_S2B**), consistent with a faster repopulation of i-cells in *Ho_CR* than in *Ho_CS*. This result confirms the dramatic and irreversible impact of gametogenesis on somatic interstitial lineages in *Ho_CS* but not in *Ho_CR*. As i-cell loss can be induced at 18°C by drugs or heat-shock, we compared the profiles of i-cell markers in aging *Ho* animals and in drug-treated or heat-shocked *Hv* animals (Wenger et al. 2016). We noted for 10/20 a more drastic reduction after drugs or heat-shock, indicating that gametogenesis leads to a more partial i-cell depletion (**Figure_S2C**).

### Aging can be induced independently of gametogenesis in *Ho_CS*

To test whether i-cell depletion obtained in the absence of gametogenesis suffices to induce aging, we exposed animals maintained at 18°C to HU and left them unfed (Figure 2A). We noted in *Ho_CS* animals a typical aging phenotype as observed after cold-induced gametogenesis, although arising faster, including the epithelial myofiber disorganization (Figure 2B-2G). By contrast, we did not record any sign of aging in *Ho_CR* or *Hv* animals that rather exhibit a starvation phenotype, characterized by a reduced size but no dysmorphic features. Nevertheless *Ho_CS* as *Ho_CR* die within six weeks, while *Hv* animals

**Figure 2:**
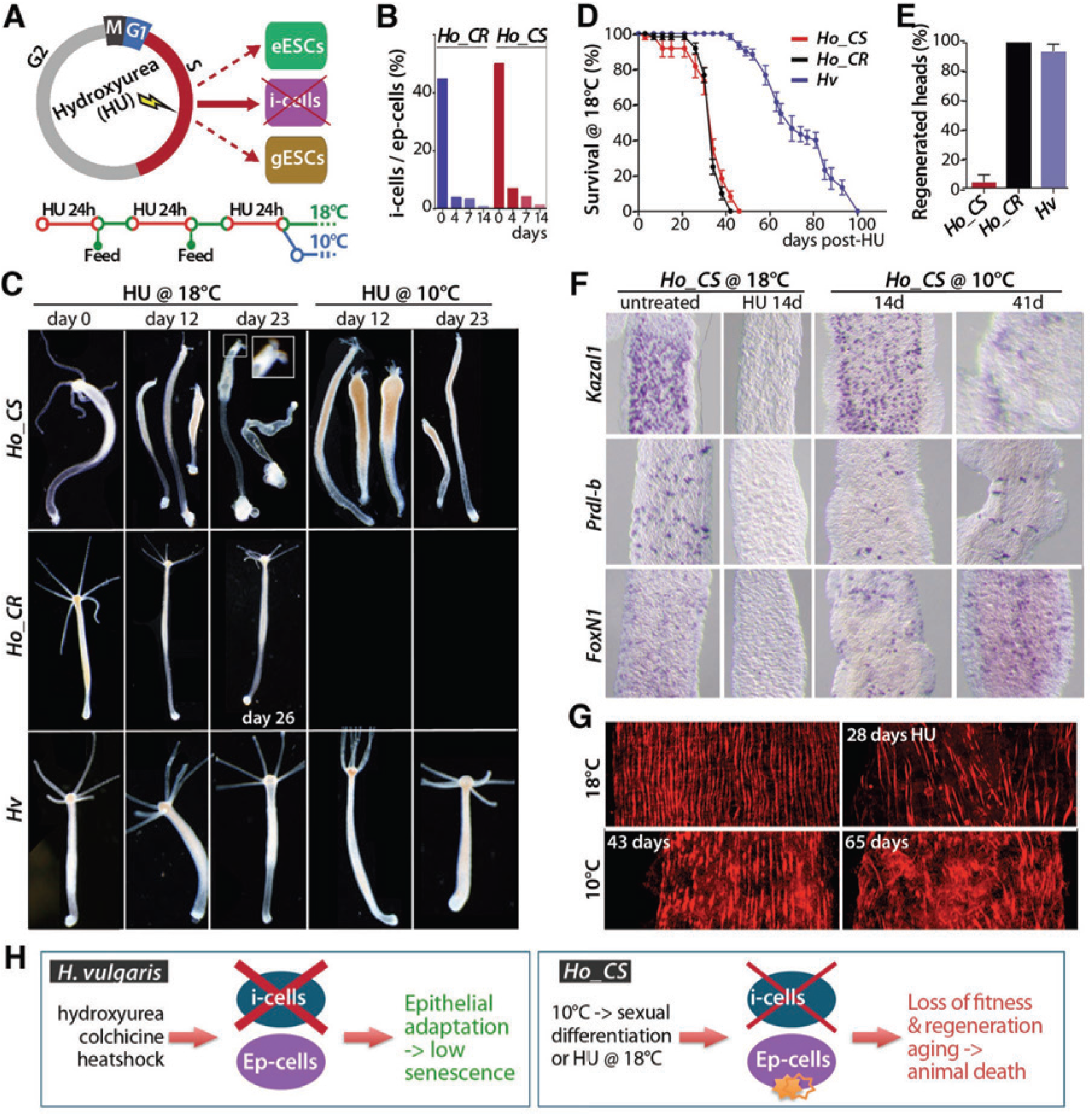
Pharmacological induction of aging in *Ho_CS* animals maintained at 18°C. **(A)** Pulse treatments of hydroxyurea (HU) eliminate all ISCs but not ESCs as i-cells cycle 3-4x faster. **(B)** Interstitial cell ratios measured in HU-treated ***Ho_CR*** and ***Ho_CS*** animals maintained at 18°C. **(C)** HU-induced morphological changes noted in ***Ho_CS, Ho_CR*** and *Hv* animals. Aging signs appear only in HU-treated ***Ho_CS*** animals. **(D)** Survival rate after HU treatment (n=6x 10). **(E)** Head regeneration rate of animals bisected 9 days after HU treatment (n=3x 10). **(F)** Expression of ***Kazal-1, prdl-b*** and ***Fox-N1*** (i-cell marker) in ***Ho_CS*** animals at indicated days after HU-treatment or transfer to 10°C. **(G)** Phalloidin staining of eESCs in HU-treated or cold-exposed ***Ho_CS*** animals. **(H)** Scheme comparing the impact of i-cell loss in *Hv* animals where ESCs adapt (Wenger et al. 2016) and in ***Ho_CS***, where a more limited loss is lethal, suggesting a lack of epithelial adaptation.

resist twice longer to the HU-induced i-cell depletion (Figure 2D). In conclusion, ISC loss in the absence of gametogenesis suffices to induce aging phenotypes in *Ho_CS* but not in *Ho_CR* animals. This result suggests that epithelial cells adapt to i-cell loss in *Ho_CR* or in *Hv* but not in *Ho_CS* (Figure 2H). Next we tested the properties of epithelial cells in aging and non-aging *Hydra*.

### Deficient autophagy inducibility in response to starvation in *Ho_CS*

We first tested the resistance to starvation of *Ho_CS, Ho_CR*, and *Hv* animals maintained at 18°C and noted the faster loss of pigmentation and progressive size decrease in *Ho* strains than in *Hv* animals that live twice longer (**Figure_S3**). However despite their similar survival rate, *Ho_CS* animals undergo a more dramatic reduction of their tissue layers than *Ho_CR*, particularly the gastroderm (Figure 3A). As starved *Hydra* rapidly induces epithelial autophagy (Buzgariu et al. 2008; Chera et al. 2009), we analyzed autophagosome formation by anti-LC3B immunostaining. The ubiquitin-like protein ATG8/LC3 is an essential component of autophagosomes (Birgisdottir et al. 2013), present as four copies in *Hydra* (LC3A/B, LC3C, GABARAPL1, GABARAPL2), all expressed at high levels in ESCs (**Figure_S4**). In 18°C-maintained animals we detected typical LC3+ vacuoles after one-day starvation (Figure 3B). After 17 starvation days their number doubles in *Hv* and *Ho_CR* animals, but not in *Ho_CS* where their size tends to enlarge (Figure 3C). In cold-maintained *Ho_CS* animals, neither the size, nor the number of autophagosomes significantly increases during aging, suggesting that autophagy regulation is deficient in *Ho_CS*. We also tested the Ulk1 inhibitor SBI-0206965 (Egan et al. 2015) that did not affect animal survival (**Figure_S3C**).

**Figure 3:**
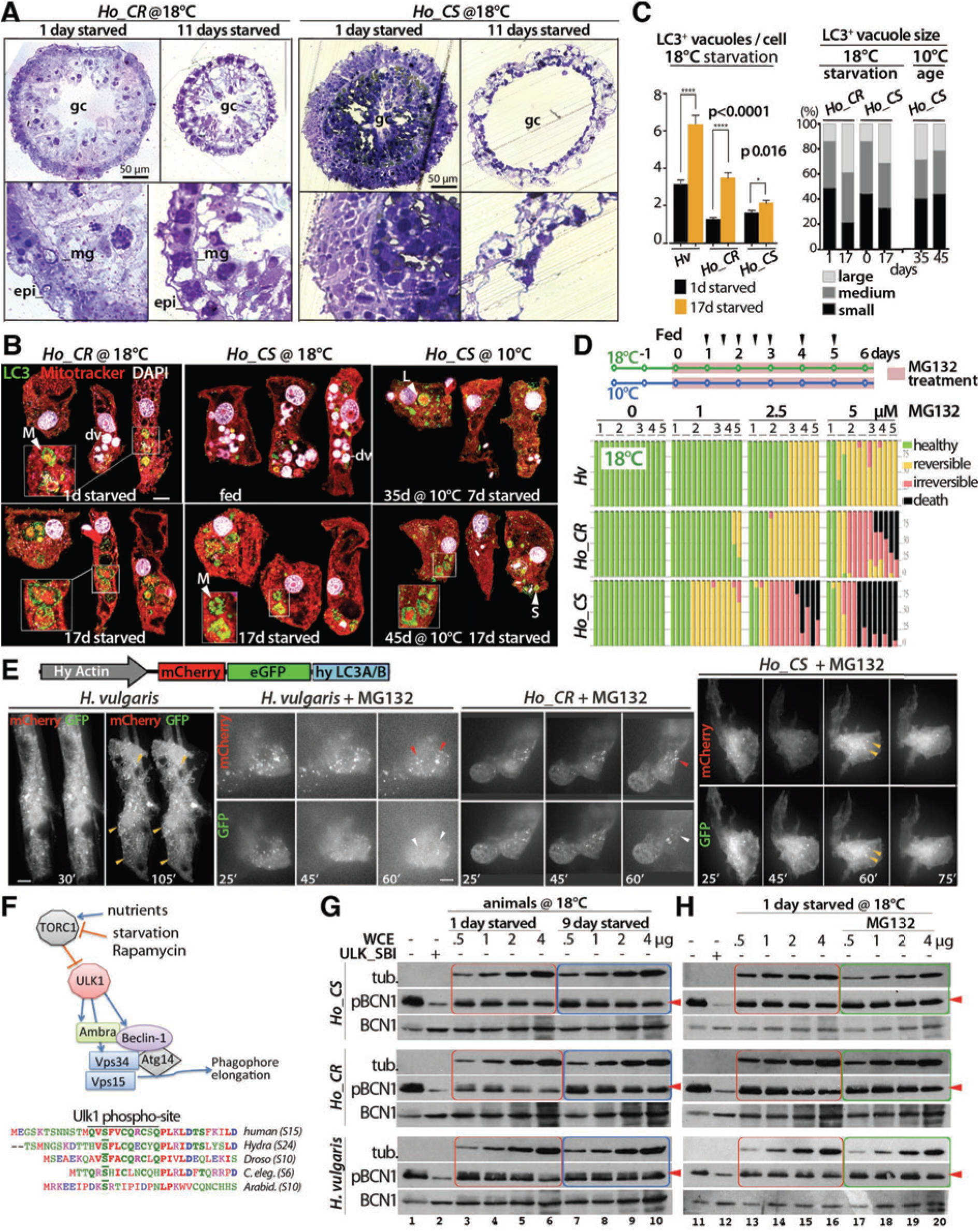
Deficient inducibility of the autophagy flux in *Ho_CS* animals. **(A)** Toluidine-stained transversal sections of gastric regions from fed and starved animals. Lower panels correspond to enlarged areas of above panels; epi: epidermis, gc: gastric cavity, mg: mesoglea. **(B)** Detection of large (L), medium (M) and small (S) autophagic vacuoles (av, arrowheads) and digestive vacuoles (dv, pink) in epithelial cells immunodetected for LC3 (green) and stained with mitotracker (red) and DAPI (white). **(C)** Number (left) and size (right) of LC3+ vacuoles counted in at least 100 ESCs per condition, p-values calculated using unpaired t-test. **(D)** Toxicity recorded in animals (n= 2x 10 /strain) maintained at 18°C (left) or 10°C (right) and continuously exposed to the proteasome inhibitor MG132 at indicated concentrations. **(E)** Live imaging of ESCs transiently expressing the mCherry-eGFP-hyLC3A/B autophagy sensor in animals maintained at 18°C either untreated or exposed from t0 onwards to MG132 5 *μ*M. **(F)** Scheme showing the TORC1 regulation of the Beclin-1 complex via Ulk1 phosphorylation and the conservation of the Ulk1 phospho-site in Beclin-1. **(G, H)** *In vitro* kinase assays testing the activity of *Hydra* extracts prepared from animals, either fed or starved **(G)**, treated or not with MG132 for 16 hours **(H)** on Ulk1-dependent phosphorylation of human Beclin-1 at Ser15.

### Deficient autophagy activation in response to proteasome inhibition in *Ho_CS*

Autophagy, which plays a crucial role in proteostasis, is up-regulated upon proteasome inhibition (Lan et al. 2015). To test the response to proteasome inhibition, we exposed animals to MG132 either continuously over six days, or as a 16 hour pulse (Figure 3D, **Figure_S3D**). In both conditions, *Ho_CS* animals rapidly show signs of toxicity, far before *Ho_CR* and *Hv* animals, suggesting a lower efficiency in compensatory autophagy. To visualize the autophagy flux, we used the mCherry-eGFP-LC3 sensor (Pankiv et al. 2007), designed to anchor mCherry and eGFP in early and mature autophagosomes, while the pH-sensitive GFP fluorescence gets lost after lysosome fusion. Animals electroporated for transient epithelial expression were live imaged two days later. In *Hv* animals, the GFP fluorescence of most mCherry-GFP-hyLC3 puncta persists in untreated animals but vanishes within 75 minutes after MG132 exposure, indicating an efficient activation of the flux (Figure 3E). We noted a similar MG132-dependent activation of the autophagy flux in *Ho_CR* but not in *Ho_CS* epithelial cells where the fluorescence is diffused and the rare mCherry-GFP-hyLC3 puncta remain stable in the presence of MG132. These *in vivo* assays confirm the poor inducibility of the autophagy flux in *Ho_CS*.

### Differential regulation of Ulk1 activity in *Hv, Ho_CR* and *Ho_CS*

Two main nutrient sensing pathways, TORC and AMPK, regulate the entry into autophagy through the Ulk1 kinase complex, which once activated, phosphorylates components of the Beclin-1 complex that initiates autophagosome formation (Figure 3F) (Russell et al. 2013). To investigate Ulk1 activity in *Hydra* extracts, we first measured *in vitro* the phosphorylation level of human Beclin-1 at Ser15, a phosphorylation site conserved in *Hydra* Beclin-1 (Russell et al. 2013; Egan et al. 2015). As Ulk1 activity was undetectable in *Hydra* extracts (not shown), we used *Hydra* extracts to monitor the activity of Ulk repressors: *Hydra* extracts produced from daily fed or 9-day starved animals maintained at 18°C were coincubated with the recombinant human Ulk1 and Beclin-1 proteins. *Hv* extracts exhibit a low repressive activity on Ulk1 activity, while, as expected, *Ho_CR* extracts from fed animals inhibit Ulk1 activity more strongly than extracts from starved ones (Figure 3G, compare red and blue frames). By contrast, extracts prepared from starved *Ho_CS* animals show a limited Ulk1 derepression when compared to extracts from fed animals. These results point to a similar constitutive repression of Ulk1 activity in both *Ho* strains when compared to *Hv*. However Ulk1 activity appears derepressed in *Ho_CR* animals submitted to starvation but not in *Ho_CS* ones. In *Hv* and *Ho* strains, a 16 hours MG132 treatment only slightly modifies Ulk1 inhibition (Figure 3H, compare red and green frames), implying that MG132 enhances autophagy via a different path.

### Differential regulation of the autophagy genetic program in *Ho_CR* and *Ho_CS*

Next we analyzed how autophagy genes respond to cold induction by analyzing in *Ho* animals maintained at 18°C or 10°C the RNA-seq expression profiles of 74 genes linked to autophagy (**Figure_S5, Figure_S6, Table-S1**). We found 50 genes specifically regulated at 10°C, 34/50 transiently up-regulated in both strains but 20/34 delayed by 10 days in *Ho_CS* when compared to *Ho_CR*, 5/50 up-regulated in *Ho_CR* but poorly in *Ho_CS (AMBRA1, ATG16L1, BECN1, RAB24, VAMP7)*, 4/50 transiently up-regulated in *Ho_CS* but poorly in *Ho_CR (ATG2B, ATG4C, PLEKHF2, TOLLIP)*, 7/50 up-regulated at late time-points in *Ho_CS (ATG4B, ATG7, CALRC, DAPK1, LAMP1, NBR1, p62/SQSTM1)*. These profiles suggest deficiencies at several levels of the autophagy flux in *Ho_CS* when compared to *Ho_CR*. The lack of regulation in *Ho_CS* of *Ambra1* and *Beclin-1* points to a deficient initiation of phagosome formation after cold transfer, while the stronger up-regulation in *Ho_CS* of *Ulk1/2* and *Atg13*, two components of the Ulk1 complex crucial for Beclin-1 complex activation, possibly reveals a regulatory feedback loop mechanism. In agreement with a deficient initiation, we noted the delayed up-regulation of *ATG4B*, encoding a protease involved in LC3 cleavage and maturation, essential for autophagosome elongation. Also deficient in *Ho_CS*, the upregulation of *Rab24*, a small GTPase required for terminating starvation-independent autophagy (Yla-Anttila et al. 2015). Finally the late upregulation in *Ho_CS* of *NBR1* and *p62/SQSTM1* that encode autophagic cargo receptors (Bjorkoy et al. 2005; Johansen and Lamark 2011), likely reflects a blockade of the autophagy flux. Indeed p62/SQSTM1 is an evolutionarily-conserved protein that accumulates when its cargo is not properly degraded. In *Hydra p62/SQSTM1* is ubiquitously expressed, predominantly in epithelial cells, steadily accumulating during aging (Figure_S7), providing an additional evidence for an inefficient autophagic flux in *Ho_CS*.

### Anti-aging effect of Rapamycin in *Ho_CS*

Rapamycin acts as a potent inhibitor of MTORC1 (Mechanistic Target of Rapamycin Complex 1), a complex that prevents autophagy. Consequently, Rapamycin enhances autophagy, thus promoting resistance to starvation (Shen and Mizushima 2014) and prolongs lifespan in multiple species, including yeast and mice (Fontana et al. 2010). To test the effect of Rapamycin treatment on aging *Ho_CS*, we continuously exposed the animals to Rapamycin from day-2 after transfer to 10°C. We noted a prolonged maintenance of animal fitness as observed at day-58 (Figure 4A), a time when *Ho_CS* animals are severely degenerated with no visible head structures, while most Rapamycin-treated ones still harbor tentacles, contract and survive several weeks longer (Figure 4B). Also Rapamycin treatment dramatically improves regeneration in aging *Ho_CS* animals (**Figure_S9A**). Thus Rapamycin provides sustained antiaging benefits in *Ho_CS* animals. Nevertheless Rapamyin changes testis morphology in both strains, with testes exhibiting a flatten shape and containing fewer mature sperm cells (Figure 4C, **Figure_S9B**).

**Figure 4:**
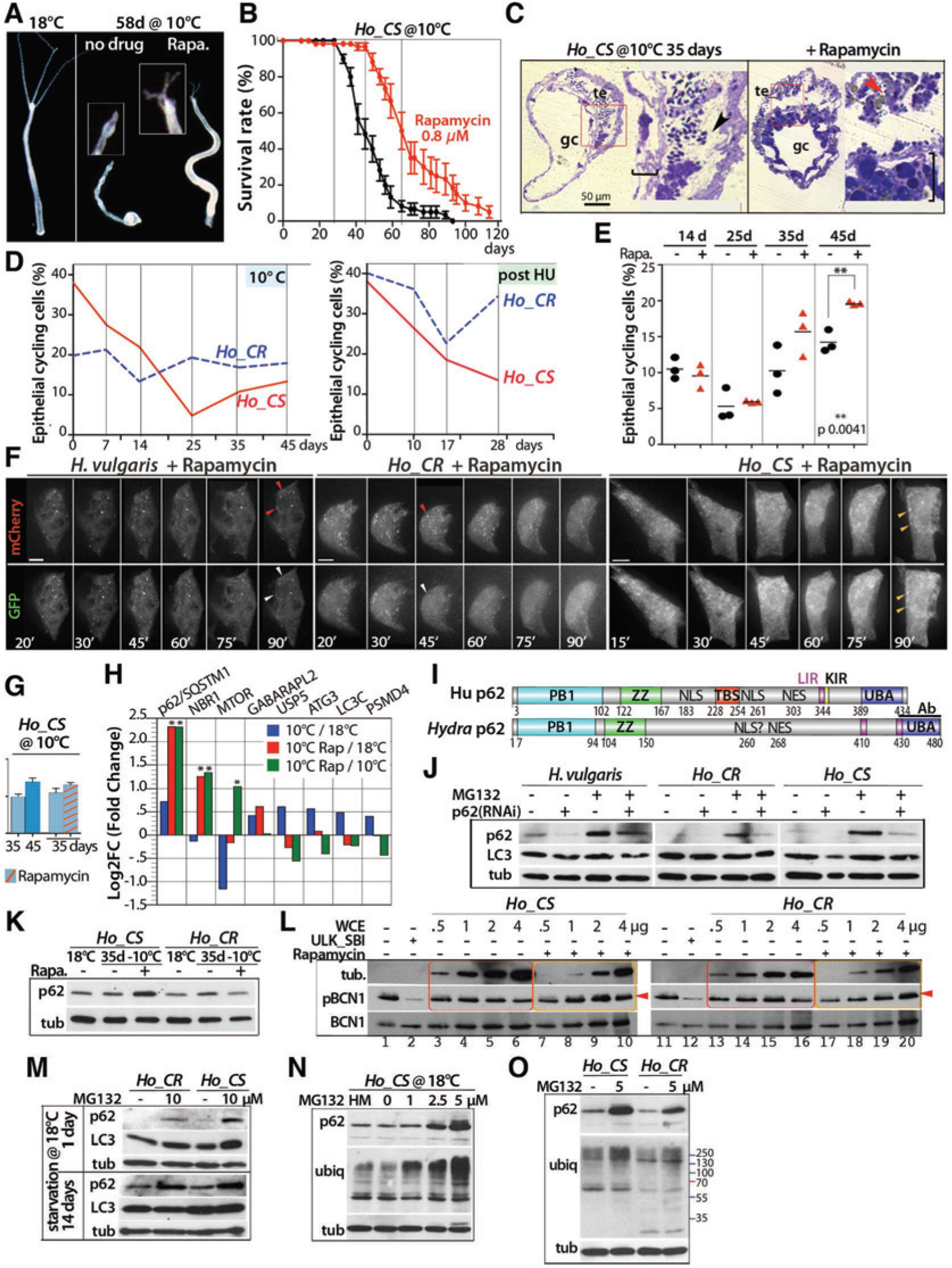
Rapamycin treatment delays aging in *Ho_CS* without enhancing the autophagy flux. **(A)** Aging morphological phenotype and **(B)** survival rates measured in *Ho_CS* animals exposed or not to Rapamycin from day-3 at 10°C. **(C)** Toluidine-stained transversal sections of gastric regions from *Ho_CS* animals maintained at 10°C for 35 days exposed or not to Rapamycin. Red squares: enlarged areas shown on the right of each panel; gc: gastric cavity, te: testis; brackets: thickness of the gastroderm; black arrowhead: sperm cells in the testis lumen; red arrowhead: sperm cells engulfed in an epithelial cell. **(D, E)** BrdU-labeling index values measured after 96 hours BrdU exposure (BLI-96) in animals maintained at 10°C (D, left), HU-treated (D, right), or in *Ho_CS* treated or not with Rapamycin (E). In D, different BLI values at t0 are due to differences in the feeding diet the two previous weeks *(Ho_CS:* 4x/week, *Ho_CR:* 2x/week). **(F)** Live imaging of epithelial cells transiently expressing the mCherry-eGFP-hyLC3A/B autophagy sensor in animals maintained at 18°C and exposed to Rapamycin (0.8 *μ*M) from t0. **(G)** Number of LC3-positive vacuoles counted in at least 100 ESCs of *Ho_CS* maintained at 10°C treated or untreated with Rapamycin, p-values calculated using unpaired t-test. **(H)** Proteomic analysis performed on *Ho_CS* animals maintained for 35 days at 18°C or at 10°C exposed or not to Rapamycin for 32 days, *: 0.05, **: 0.001 significance. **(I)** Structure of the human and *Ho* p62/SQSTM1 (p62) proteins (see alignment in **Figure_S8**). The black bar indicates the region used to raise an anti-Hydra p62 antibody. **(J)** p62 levels in animals RNAi-silenced for p62 and exposed or not to MG132 for 16 hours; tub: α-tubulin. **(K)** p62 levels in animals treated or not with Rapamycin. **(L)** *In vitro* kinase assays testing the activity of *Hydra* extracts prepared from animals treated or not with Rapamycin for 13 days on Ulk1-dependent phosphorylation of human Beclin-1. **(M)** p62 and LC3 levels in animals starved for 14 days or not, and exposed to MG132 or not for 16 hours. **(N, O)** p62 levels and ubiquitin patterns in animals maintained at 18°C and exposed to MG132 or not for 16 hours at indicated conditions.

Next we tested whether Rapamycin promotes epithelial proliferation as on histological sections, we noted a thicker gastroderm in Rapamycin-treated animals (Figure 4C). During cold exposure, cycling activity remains rather stable in *Ho_CR* animals but progressively declines in *Ho_CS* ones, from 40% cycling cells at day-0 to 5% at day-25, although slightly recovering later (Figure 4D). Similarly after HU treatment at 18°C, cell cycling remains roughly stable in *Ho_CR* although transiently reduced at day-17, while becoming irreversibly reduced in *Ho_CS* animals. We also noted a delayed activation of a large subset of cell cycle genes in *Ho_CS* compared to *Ho_CR* (not shown), in agreement with the inability of *Ho_CS* animals to restore and maintain epithelial proliferation after i-cell loss. In Rapamycin-treated *Ho_CS* animals maintained at 10°C, stem cell proliferation also dramatically declines but is rescued at day-35 and day-45 (Figure 4E). Hence Rapamycin might delay aging by restoring epithelial proliferation after i-cell loss.

### Rapamycin does not enhance the autophagy flux in *Ho_CS*

Next we tested the regulation of the autophagy flux in *Ho_CS* animals exposed to Rapamycin. The *in vivo* analysis showed, as with MG132, a rapid flux activation after Rapamycin exposure in *Hv* and *Ho_CR* but not in *Ho_CS* cells, where we found the mCherry-eGFP-LC3A/B fluorescence predominantly cytoplasmic and the rare mCherry+/GFP+ puncta persisting after Rapamycin exposure (Figure 4F). Accordingly the number of LC3-positive vacuoles is not modified upon Rapamycin (Figure 4G). By quantitative proteomics
we also detected p62/SQSTM1 up-regulated 1.6x in 35-day old *Ho_CS* animals and 5.6x in Rapamycin-treated ones (Figure 4H). We raised an antibody against the p62/SQTSM1 C-terminus (Figure 4I, **Figure_S8**), validated its specificity on extracts from animals silenced for p62/SQSTM1 (Figure 4J), confirmed its up-regulation when aging and Rapamycin are combined (Figure 4K) and showed its colocalization with LC3+ autophagosomes (**Figure_S9d, Movie S3**). The Ulk1 kinase assays testing extracts from animals maintained at 10°C for 13 days, exposed or not to Rapamycin, show a derepression of Ulk1 activity by *Ho-CR* extracts, and a partial one by *Ho-CS* extracts (Figure 4L). This suggests that Rapamycin only partially counteracts the Ulk1 repressor identified in *Ho_CS* extracts. In animals maintained at 18°C, starvation and MG132 exposure also lead to p62/SQSTM1 accumulation, higher in *Ho_CS* than in *Ho_CR* (Figure 4M), together with an accumulation of polyubiquitinated proteins, constitutively more abundant in *Ho_CS* than in *Ho_CR* (Figure 4N,4O, **Figure_S9C**). Hence, the enhanced p62/SQSTM1 accumulation in *Ho_CS*, whatever the autophagy inducer(s), confirms that the autophagy flux remains deficient in Rapamycin-exposed *Ho_CS* animals.

### Rapamycin promotes epithelial phagocytosis and lipid droplet accumulation

To further investigate how Rapamycin delays aging in *Hydra*, we analyzed tissue sections and noticed that epithelial phagocytosis is strongly enhanced upon Rapamycin (Figure 5). In regularly-fed animals maintained at 18°C, epithelial cells contain a dense cytoplasm, numerous mitochondria, few intracellular spaces, and numerous digestive vacuoles. After 11 starvation days, mitochondria become sparse while numerous intracellular spaces appear, as well as autophagosomes in *Hv* animals, i.e. double-membrane vacuoles with a heterogeneous content; such vacuoles are rare in *Ho_CS* animals (Figure 5A). After 35 days at 10°C, cells from Rapamycin-treated *Ho_CS* animals display a reduced cytoplasm, large intracellular spaces and sparse autophagosomes, but numerous electron lucent vacuoles resembling lipid droplets, as confirmed by Bodipy 493/503 labeling (Figure 5B). We also identified numerous sperm cells in the epithelial intracellular spaces, internalized in the cytoplasm and finally digested (Figure 5C, 5D). Rapamycin-induced epithelial phagocytosis is also visible in *Ho_CR* animals, with intracellular germ cells visible besides the digestive vacuoles (Figure 5D, 5E). These phagocytized cells, numerous at day-35, are frequently surrounded by a thick p62+ rim, often co-localizing with LC3 (Figure 5E, **Figure_S9D-H, Movie-S2**), already detected at day-9 post-transfer (**Figure_S9G**). At day-9, the Rapamycin-enhanced epithelial phagocytosis is not restricted to germ cells as numerous phagocytized small cells, typically nematoblasts or nematocytes can be detected (**Figure_S9G**). The quantification of this engulfment process shows a Rapamycin-induced phagocytosis maximal at day-36, markedly higher in *Ho_CS* than in *Ho_CR* (Figure 5G, **Figure_S9E**). For animals maintained at 18°C, the chronic exposure to Rapamycin was too toxic to detect any effect (not shown). All together, these results indicate that Rapamycin dramatically enhances the phagocytic behavior of epithelial cells in both *Ho_CS* and *Ho_CR*, likely providing a source of exogenous nutrients that promote animal survival in *Ho_CS*.

**Figure 5:**
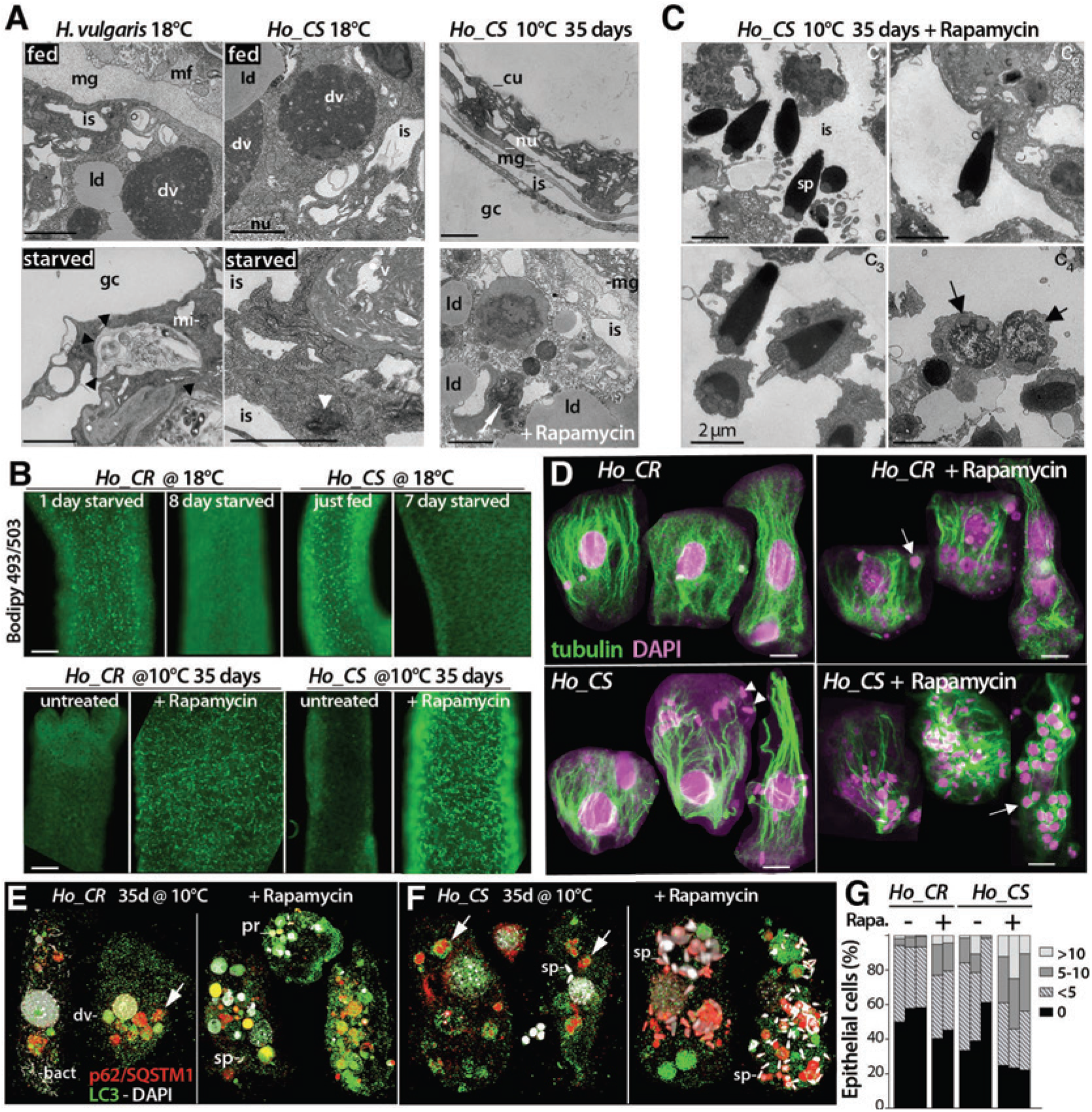
Rapamycin promotes epithelial phagocytosis and lipid droplet formation in *Hydra*. **(A)** TEM views of body column sections from *Hv* and *Ho_CS* animals maintained as indicated. Black arrowheads: autophagosome, white arrowhead: aggregate, white arrow: engulfed corpse. **(B)** Detection of lipid droplets in live animals stained with Bodipy 493/503. **(C)** Sperm cells (sp) engulfed in epithelial cells from Rapamycin-treated *Ho_CS* animals maintained at 10°C for 35 days, detected in the intracellular space (is, c1, c2), surrounded by cytoplasm (c3) and digested (c4). Black arrows: mitochondria at the base of sperm cells, white arrowheads: sperm nuclei. **(D)** Engulfed cells detected with an anti a-tubulin antibody (green) and DAPI staining (pink) in *Ho_CS* and *Ho_CR* epithelial cells at day-36 after cold transfer. Arrows point to immature germ cells and arrowheads to sperm cells. **(E, F)** Engulfed cells immuno-detected with the anti-Hydra p62/SQSTM1 (red) and the anti-human LC3 (green) antibodies, co-stained with DAPI (white). Arrows point to p62-labeled granules associated or not with LC3. Abbreviations: cu: cuticle, dv: digestive vacuole, gc: gastric cavity, is: intracellular space, ld: lipid droplets, mf: myofibril, mg: mesoglea, mi: mitochondria, nu: nucleus, pr: progenitors, sp: sperm cells. Scale bars = 2 (a,b), 10 (d,e,f), 100 (c) *μ*m. **(G)** Number of engulfed cells per epithelial cell at day-36 after cold transfer, counted in four categories: >10, 5-10, <5, 0 per cell. Each condition in triplicates (117-249 cells per sample).

### Efficient autophagy is required for fitness and survival in *H. vulgaris*

To test whether a blockade of the autophagy flux promotes aging in non-aging *Hv* animals, we disrupted autophagosome formation by knocking-down WIPI2, which recruits the Atg12–5-16L1 complex, critical for LC3 conjugation (Figure 6A) (Dooley et al. 2014). *Hydra* WIPI2 possesses both the ATG16 and the PI3P binding motifs necessary for its function (Figure 6B, **Figure_S10**). To silence *WIPI2*, we repeatedly electroporated (EP) *Hv* polyps with siRNAs (Figure 6C) and noted after EP-6 a 80% reduction in *WIPI2* transcript level (Figure 6D). Consistent with a deficient autophagic flux, we found the p62/SQSTM1 protein level twice higher in WIPI2(RNAi) animals (Figure 6D). We also noted a lower number of LC3 puncta in epithelial cells of WIPI2(RNAi) animals, where LC3 signals are diffuse and cytoplasmic even in the presence of MG132 (Figure 6E, 6F), similar to the LC3 pattern observed in *Ho_CS* epithelial cells. In MG132-treated cells of control(RNAi) animals, the number of LC3 puncta is quite stable, consistently with a robust induction of autophagy (Figure 6F). At the phenotypic level, we noted the size decrease of WIPI2(RNAi) animals after EP-7, and their apical disorganization (Figure 6G). About 60% of these animals died within the two following weeks, while no mortality was observed in control(RNAi) animals (Figure 6H). Control(RNAi) animals bisected at mid-gastric level after EP-6 regenerated normally their head, while about 30% animals WIPI2(RNAi) animals show a delayed or blocked regeneration (Figure 6I). Moreover, the WIPI2(RNAi) animals having regenerated their head rapidly died, proving their lack of fitness (not shown). In summary, knocking-down *WIPI2* suffices to block autophagy by impairing the loading of LC3 onto phagophores, resulting in the observed switch from punctuated to diffuse LC3 pattern. A prolonged autophagy blockade in the slow-aging *Hv* animals leads to a rapid loss of animal fitness, followed by animal death, mimicking the aging phenotype observed in *Ho_CS* animals.

**Figure 6:**
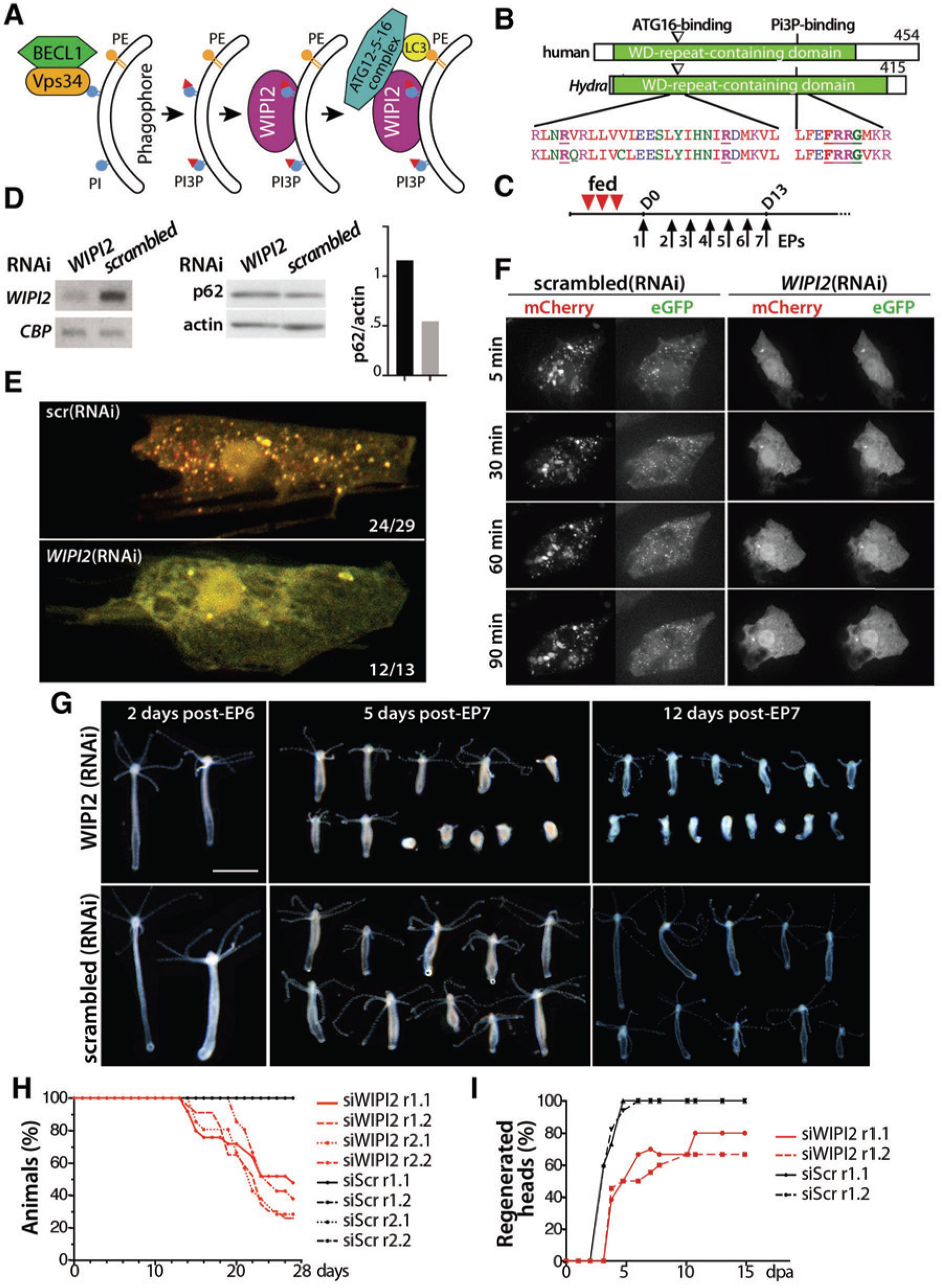
Impact of *WIPI2* silencing on autophagic flux, survival, and regeneration in *H. vulgaris*. **(A)** Scheme showing the role of WIPI2 in autophagosome formation(Dooley et al. 2014). **(B)** Structure of the human and *Hydra* WIPI2 proteins and alignment of the regions involved in ATG12-5-16 complex and PI3P binding. **(C)** RNAi silencing procedure performed on intact well-fed animals. Red arrowheads: feedings, black arrows: electroporation (EP). **(D)** *WIPI2* and ***CBP*** (CREB-binding protein) RNA levels (left) and p62 and actin protein levels (right) in *Hv* animals electroporated 6x with *WIPI2-* or *scrambled*-siRNAs. **(E, F)** LC3 pattern detected in epithelial cells expressing the mCherry-eGFP-hyLC3 autophagy sensor two days after EP7 in *WIPI2*(RNAi) or *scrambled*(RNAi) animals. Note in (e) the lack of LC3 puncta in *WIPI2(RNAi)* cells. In (f) animals were exposed to MG132 at t0. **(G-I)** Phenotypic analysis of *WIPI2(RNAi)* and scrambled(RNAi) animals: morphology after EP6 or EP7 **(G)**; mortality rates recorded after EP7 (**H**); head regeneration after bisection performed at mid-gastric level 24 hours post-EP6 (**I**). Scale bar = 1mm.

## DISCUSSION

### Epithelial adaptability as a common basis for low senescence

The aging phenotype in *Ho_CS* animals share features with mammalian aging, like loss of somatic stem cells, deterioration of the muscular network, alterations of the nervous system, behavioral alterations, global loss of health. This aging process results from epithelial deficiencies that prevent ESCs to adapt to environmental changes, rather than from direct negative effects of these changes (Figure 7). By contrast, *Hv* animals survive weeks of starvation, a period when their epithelial cells enhance their phagocytic behavior (Bosch and David 1984), activate an efficient autophagy program (Buzgariu et al. 2008; Chera et al. 2009) and reversibly slow-down cycling (Otto and Campbell 1977). Also *Hv* animals can become “epithelial”, surviving for years (if manually fed) the elimination of their i-cells and nervous system (Marcum and Campbell 1978; Sugiyama and Fujisawa 1978), as their ESCs adapt their genetic program (Wenger et al. 2016). Therefore a robust epithelial adaptability to environmental conditions evidenced by changes in cell cycle, autophagy flux, gene expression, provides a necessary and sufficient framework for low senescence in *H. vulgaris* strains (Schenkelaars et al. 2018).

**Figure 7:**
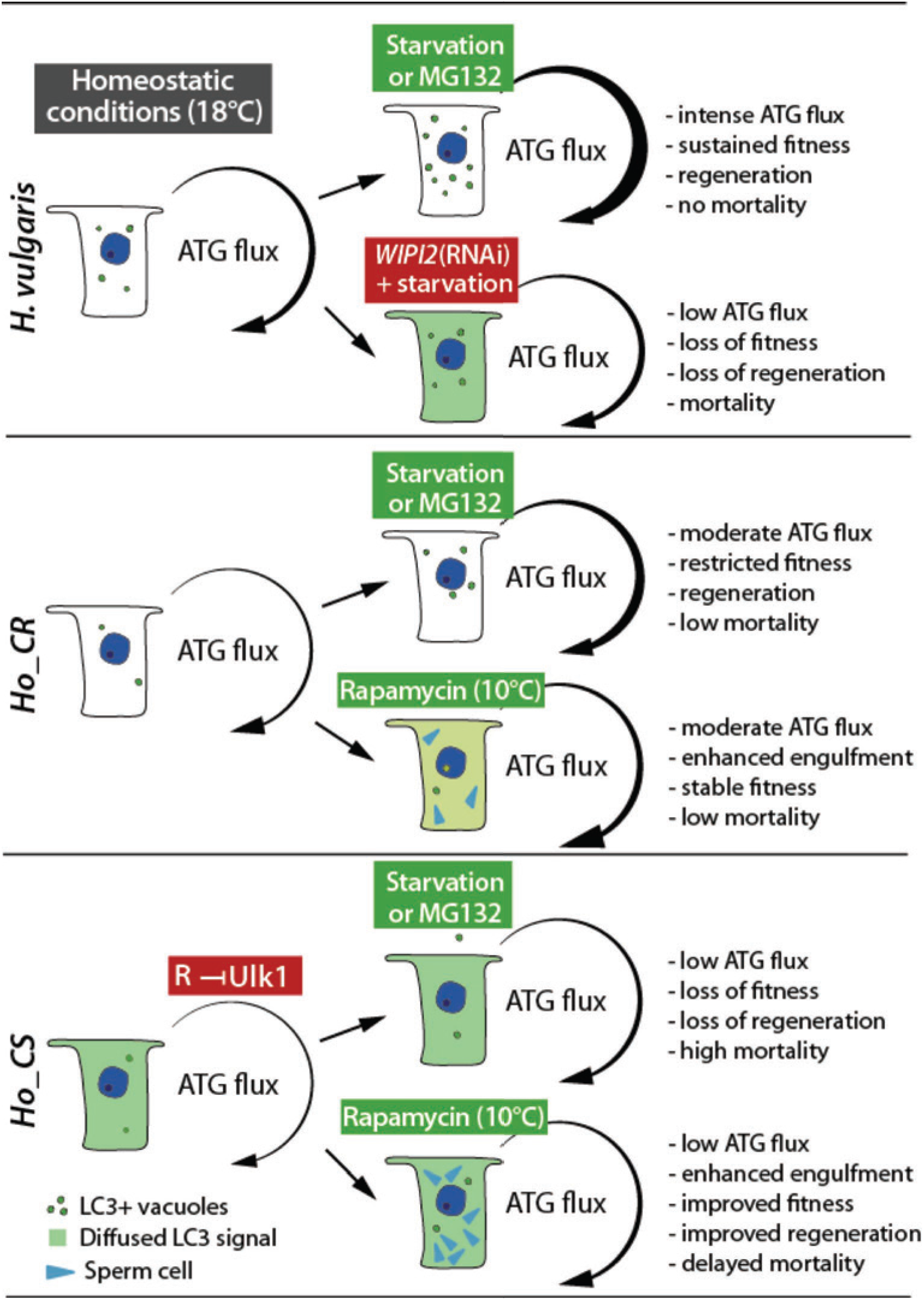
Summary view of the responsiveness of the autophagic flux in aging and non-aging *Hydra*. The responsiveness of the autophagy flux to starvation, MG132 or Rapamycin treatments was deduced from the formation of LC3-positive vacuoles, the accumulation of p62/SQSTM1 and the *in vivo* imaging of LC3 vacuoles in *Hydra* epithelial cells.

### Deficient response to proteotoxic stress in *Hydra oligactis*

The maintenance of proteostasis is critical for longevity (Morimoto and Cuervo 2009; Kaushik and Cuervo 2015). In *Hydra* we identified three distinct levels of proteostasis efficiency: low in *Ho_CS* animals that are highly sensitive to proteasome inhibition, intermediate in *Ho_CR*, and high in *Hv*. As the ubiquitin-proteasome and the lysosome-autophagosome are the two main systems that maintain proteome homeostasis (Pandey et al. 2007; Rubinsztein 2007; Lan et al. 2015), this observed difference in MG132 sensitivity between aging and non-aging *Hydra* strains supports the finding that deficient autophagy contributes to aging in *Ho_CS*. Also the molecular chaperone Hsp70, involved in protein folding, might explain the observed difference in proteostasis efficiency between *Hv* and *Ho* strains. Indeed, *Hsp70* expression is up-regulated in both strains upon heatshock but in *Ho*, transcripts are unstable and the protein not efficiently translated (Gellner et al. 1992; Brennecke et al. 1998). Thus as in yeast, *Drosophila, C. elegans* or mammals (Cuervo 2008) an efficient autophagy appears necessary to prevent aging in *Hydra*.

### Efficient epithelial autophagy is required for maintaining epithelial cell cycling in *Hydra*

In *Hv* animals submitted to physiological starvation, numerous interstitial cells undergo apoptosis and are phagocytosed by the epithelial cells, a process supposed to sustain cell proliferation and animal survival (Bosch and David 1984). Here we identified a similar mechanism, where the autophagy deficiency identified in *Ho_CS* primarily causes aging, by reducing epithelial proliferation to a point where regeneration and animal survival are compromised. *Ho_CR* but not *Ho_CS* animals adapt and ultimately survive, as they induce an efficient autophagy that allows a sustained epithelial cell cycling. Three arguments support this scenario, (i) the correlation between epithelial self-renewal and autophagy levels observed in *Ho_CS* and *Ho_CR* animals that undergo gametogenesis at 10°C; (ii) the rescue of epithelial cell cycling when Rapamycin-induced phagocytosis provides a source of nutrients; (iii) the aging process induced in *Hv* animals upon blockade of autophagy. Therefore autophagy seems to protect against aging by maintaining stem cell renewal. Similarly, in aging mammals, autophagy is crucial for the survival of hematopoietic stem cells (Warr et al. 2013; Gomez-Puerto et al. 2016), for the maintenance of muscle satellite cells (Garcia-Prat et al. 2016).

### The Rapamycin-induced engulfment behavior as a novel modulator of aging

MTORC1 suppresses autophagy through several mechanisms (Wullschleger et al. 2006; Shen and Mizushima 2014) and its inhibition via the macrolide Rapamycin de-represses autophagy (Hosokawa et al. 2009). In aging *Ho_CS* animals, Rapamycin does not rescue the autophagy flux but rather promotes an enhanced engulfment behavior of the epithelial ectodermal cells, which digest the surrounding small cells including germ cells, a behavior rarely observed in untreated cells. This engulfment process resembles *entosis* in which live cells become engulfed within single-membrane LC3-associated vesicles (Florey et al. 2011) and then degraded thanks to the support of MTORC1 (Krajcovic et al. 2013), but also *LC3-associated phagocytosis (LAP)*, where LC3 associates with single-membrane phagosomes containing pathogens (Sanjuan et al. 2007). A third possibility would be *phagoptosis* where viable cells that display “eat-me” signals or lose the “don’t-eat-me” signals can be phagocytized (Brown and Neher 2012). The Rapamycin-induced increase in p62/SQSTM1 levels in aging animals and the thick p62/SQSTM positive rim surrounding most engulfed cells suggest that p62/SQSTM1 is involved in this engulfment process. Unfortunately the *p62/SQSTM1* (RNAi) knock-down was too partial to prevent the Rapamycin-induced accumulation of p62/SQSTM1 (Figure 4J). This beneficial Rapamycin-induced phagocytic behavior of epithelial cells is a new finding with no equivalent in any other system to our knowledge. If artificially promoted, it could rescue an active metabolism in cells where autophagy is deficient and thus act as a potent anti-aging mechanism. However Rapamycin may also delay the aging phenotype by reducing the amount of energy allocated to the germline.

### Complex regulation of Rapamycin-induced lipid droplet accumulation in *Ho_CS*

Rapamycin treatment also leads to the production of abundant lipid droplets in *Hydra* epithelial cells, likely as a result of the nutrient inflow from the phagocytosed cells and/or by changes in lipid metabolism caused by MTORC1 inhibition (Thiam et al. 2013). In yeast, TORC1 inhibition promotes LD formation (Madeira et al. 2015) similarly to what is observed in *Hydra*. In addition, a chronic exposure to Rapamycin prevents the assembly of the TORC2 complex (Sarbassov et al. 2006), possibly modulating the lipid metabolism via TORC2. In *C. elegans*, knocking-down Rictor enhances the deposition of lipid storage via LD formation (Jones et al. 2009). More generally, Rapamycin increases the level of triacyl-glycerides as observed in *Drosophila* (Bjedov et al. 2010). As an alternative mechanism contributing to LD accumulation, the deficient autophagy characterized in *Ho_CS* might affect lipophagy, a selective type of autophagy that uses most components of the macroautophagy machinery. Hence several distinct mechanisms might contribute to the observed LD accumulation in Rapamycin-treated *Ho_CS* animals.

In summary, this study highlights the value of *Hydra* as a novel model system for aging studies, showing the impact of autophagy on the maintenance of an active stock of stem cells, and the beneficial anti-aging effect of daily exposure to Rapamycin including the maintenance of regeneration. These Rapamycin effects appear associated with energy recovery through digestion of engulfed cells, modulations of lipid metabolism, and proteostasis maintenance possibly via TORC2.

## SUPPLEMENTAL FILE

Methods, 10 supplemental Figures, one Table and four Movies are available in the supplemental file.

## ACKNOWLEDGMENTS

This research was supported by the National Institute of Health (grant R01AG037962), the Swiss National Science Foundation (SNF 31003A_149630, 31003_169930), the canton of Geneva. The authors thank H. Shimizu (NIG Mishima, Japan) who provided *Ho_CS* polyps, D. Chiappe and R. Hamelin from Proteomics Core Facility at EPFL, D. Benoni, C. Perruchoud and M-L. Curchod for excellent technical support, A-M. Cuervo, R. Loewith and T. Soldati for discussions and useful advices, R. Loewith and T. Lamark for helpful comments on the manuscript.

## AUTHOR CONTRIBUTIONS

B. G., S.T., Q.S. and Y.W. designed the experiments and validated the results; S.T., Q.S., N.S., K.E., K.F., W.B. and C. B. performed the experiments; Y.W. and S.T. curated the transcriptomic and proteomic data; S.T., Q.S. produced the original draft, revised by S.A. and B.G. Funding was obtained by S.A. and B.G. B.G. supervised this study.

